# Robust Pavlovian-to-Instrumental and Pavlovian-to-Metacognitive Transfers in human reinforcement learning

**DOI:** 10.1101/593368

**Authors:** Chih-Chung Ting, Stefano Palminteri, Jan B. Engelmann, Maël Lebreton

## Abstract

In simple instrumental-learning tasks, humans learn to seek gains and to avoid losses equally well. Yet, two effects of valence are observed. First, decisions in loss-contexts are slower, which is consistent with the Pavlovian-instrumental transfer (PIT) hypothesis. Second, loss contexts decrease individuals’ confidence in their choices – a bias akin to a Pavlovian-to-metacognitive transfer (PMT). Whether these two effects are two manifestations of a single mechanism or whether they can be partially dissociated is unknown. Here, across six experiments, we attempted to disrupt the PIT effects by manipulating the mapping between decisions and actions and imposing constraints on response times (RTs). Our goal was to assess the presence of the metacognitive bias in the absence of the RT bias. Were observed both PIT and PMT despite our disruption attempts, establishing that the effects of valence on motor and metacognitive responses are very robust and replicable. Nonetheless, within- and between-individual inferences reveal that the confidence bias resists the disruption of the RT bias. Therefore, although concomitant in most cases, PMT and PIT seem to be – partly – dissociable. These results highlight new important mechanistic constraints that should be incorporated in learning models to jointly explain choice, reaction times and confidence.

## Introduction

In the reinforcement learning context, reward-seeking and punishment-avoidance present an intrinsic and fundamental informational asymmetry. In the former situation, accurate choice (i.e., reward maximization) *increases* the frequency of the reinforcer (the reward). In the latter situation, accurate choice (i.e., successful avoidance), optimal behavior *decreases* the frequency of the response. Accordingly, any simple incremental “law-of-effect”-like model, would predict higher performance in the reward seeking compared the punishment avoidance situation. Yet, humans learn to seek reward and to avoid punishment equally-well (Fontanesi et al., 2019; Guitart-Masip et al., 2012; Palminteri et al., 2015). This is not only robustly demonstrated in experimental data, but also nicely explained by context-dependent reinforcement-learning models (Fontanesi et al., 2019; Palminteri et al., 2015), which can be seen as formal computational instantiation of Mowrer’s two-factor theory (Mowrer, 1952). On top of this remarkable symmetry in choice accuracy between gain and loss contexts, two sets of recent studies independently reported that outcome valence asymmetrically affects confidence and response times (RTs). First, learning from punishment increases individuals’ RTs, slowing down the motor execution of the choice (Fontanesi et al., 2019; Jahfari et al., 2019). This robust phenomenon is consistent with the Pavlovian-to-Instrumental Transfer (PIT) hypothesis, which posits that desirable contexts favor motor execution and approach behavior, while undesirable contexts hinder them (Boureau and Dayan, 2011; Guitart-Masip et al., 2012). Second, learning from punishment decreases individuals’ confidence in their choices (Lebreton et al., 2019). The confidence judgement is a metacognitive operation defined as the subjective estimation of the probability of being correct (Fleming and Daw, 2017; Pouget et al., 2016; Yeung and Summerfield, 2012). The demonstrations that confidence judgments can be biased by the outcome valence in different tasks (Lebreton et al., 2018, 2019) suggest that, similarly to instrumental processes, metacognitive processes could also be under the influence of Pavlovian processes – a phenomenon that we refer to as Pavlovian-to-Metacognition Transfer (PMT).

Here, we aimed to investigate the link between PIT and PMT. We focused on two research questions: first, are PIT and PMT robust and replicable? Second, can PMT be observed in the absence of PIT? Regarding the second question, previous research has yielded conflicting results that generated two opposing predictions. On the one hand, numerous studies documented behavioral and neural dissociations between perceptual, cognitive or motor operations, and confidence or metacognitive judgments (Fleming et al., 2012; Miele et al., 2011; Qiu et al., 2018). Likewise, brain lesions and stimulation protocols have been shown to disrupt confidence ratings and metacognitive abilities without impairing cognitive or motor functions (Fleming et al., 2014, 2015; Rounis et al., 2010) - although see also (Bor et al., 2017). These dissociations between decision and metacognitive variables suggest that the confidence bias (PMT) could be observed in the absence of a response time bias (PIT).

On the other hand, several studies suggest that decision and metacognitive variables are tightly linked– both in perceptual (Geller and Whitman, 1973; Vickers et al., 1985) and value-based tasks (De Martino et al., 2013; Folke et al., 2016; Lebreton et al., 2015). This coupling is notably embedded in many sequential-sampling models which rely on a single mechanism to produce decisions, response times and confidence judgments (van den Berg et al., 2016; De Martino et al., 2013; Moran et al., 2015; Pleskac and Busemeyer, 2010; Ratcliff and Starns, 2009, 2013; Yu et al., 2015). Beyond this mechanistic hypothesis, it was also recently suggested that people use their own RT as a proxy for stimulus strength and certainty judgments, creating a direct, causal link from RT to confidence (Desender et al., 2017; Kiani et al., 2014). These results could imply that our previously reported effects of valence on confidence – the so called PMT (Lebreton et al., 2019) - are no more than a spurious consequence of the effect of valence on RTs – PIT (Fontanesi et al., 2019; Jahfari et al., 2019). In other words, participants could have simply observed that they were slower in the loss context, and used this information to generate lower confidence judgments in these contexts.

In order to address our research questions, we developed several versions of a probabilistic, instrumental-learning task, where participants have to learn to seek rewards or to avoid losses (Fontanesi et al., 2019; Lebreton et al., 2019; Palminteri et al., 2015). We attempted to cancel the effects of losses on RTs (i.e. the PIT) while recording confidence judgments to assess the presence of the PMT. To this end, we modified the standard mapping between the available options and the way participants could select them, thereby disrupting the link between decision and motor execution of the choice. In another experiment, we also used a different strategy, and imposed time pressure on the choice to constrain decision time.

In total, we used two published datasets (Lebreton et al., 2019) and original data collected from four new experiments, where we manipulated in several ways the option-action mapping (experiment 3-5) and applied time pressure (experiment 6). We then tested (1) the robustness of PIT and PMT; (2) whether the effects of PMT could be observed in the absence of PIT effects. Overall, our results show that response times are slower in loss than gain contexts in almost all experiments. In other words, the PIT is highly robust, as it survived most of our disruption attempts, despite being severely attenuated. In all datasets, confidence was lower in loss than in gain contexts, indicating that the PMT is highly replicable, and is robust to variations in PIT effect sizes. The PMT is also observed in the condition where PIT was successfully canceled, suggesting that PIT and PMT are – partly – dissociable.

## Material and Methods

### Subjects

All studies were approved by the local Ethics Committee of the Center for Research in Experimental Economics and political Decision-making (CREED), at the University of Amsterdam. All subjects gave informed consent prior to partaking in the study. The subjects were recruited from the laboratory’s participant database (www.creedexperiment.nl). A total of 108 subjects took part in this set of 6 separate experiments (see **Table 1**). They were compensated with a combination of a show-up fee (5€), and additional gains and/or losses depending on their performance during the learning task: experiment 1 had an exchange rate of 1 (in-game euros = payout); experiments 2-6 had an exchange rate of 0.3 (in game euros = 0.3 payout euros, participants were clearly informed of this exchange rate). In addition, in experiments 2-6, three trials (one per session) were randomly selected for a potential 5 euros bonus each, attributed based on the confidence incentivization scheme (see below).

**Table 1.** Demographics and behavior. The correlation between confidence and performance was performed at the session level using Pearson’s R, then averaged at the individual level. Reported statistics correspond to a random-effects analysis (one sample t-test) performed at the population level. STD: standard deviation. SEM: standard error of the mean. T: Student t-value.

|  | Exp. 1 | Exp. 2 | Exp. 3 | Exp. 4 | Exp. 5 | Exp. 6 |
| --- | --- | --- | --- | --- | --- | --- |
| **Gender**<br>M/F | 8/10 | 8/10 | 10/8 | 10/8 | 6/12 | 9/9 |
| **Age**<br>mean ± STD | 24.6 ± 8.50 | 24.6 ± 4.30 | 22.72 ± 3.24 | 23.84 ± 4.12 | 20.61 ± 1.77 | 22.35±3.49 |
| **Performance**<br>mean ± SEM | 76.50 ± 2.38 | 77.04 ± 1.69 | 80.00 ± 2.82 | 75.33 ± 2.34 | 73.40 ± 2.83 | 63.60 ± 2.88 |
| **Confidence**<br>mean ± SEM | 79.19 ± 1.49 | 81.11 ± 1.58 | 78.78 ± 2.61 | 78.35 ± 2.24 | 78.09 ± 1.75 | 72.99 ± 2.14 |
| **Correlation(conf, RT)**<br>mean ± SEM<br>t(17)<br>(P-val) | -0.30 ± 0.05<br>-5.31<br>($<0.001$)*** | -0.41 ± 0.03<br>-13.32<br>($<0.001$)*** | -0.18 ± 0.03<br>-5.55<br>($<0.001$)*** | -0.16 ± 0.03<br>-5.42<br>($<0.001$)*** | -0.10 ± 0.02<br>-4.87<br>($<0.001$)*** | -0.12 ± 0.04<br>-3.03<br>(0.008)** |

### Learning task - General

In this study, we iteratively designed six experiments, aiming at investigating the impact of context valence and information on choice accuracy, confidence and response times, in a reinforcement-learning task. All experiments were adapted from the same basic experimental paradigm (see also **Figure 1 and Figure S. 1**): participants repeatedly faced pairs of abstract symbols probabilistically associated with monetary outcomes (gains or losses), and they had to learn to choose the most advantageous symbol of each pair (also referred to as context), by trial and error. Two main factors were orthogonally manipulated (Palminteri et al., 2015): valence (i.e. some contexts only provide gains, and others losses) and information (some contexts provide information about the outcome associated with both chosen and unchosen options –complete information-while others only provided information about the chosen option –partial information). In addition, at each trial, participants reported their confidence in their choice on a graded scale as the subjective probability of having made a correct choice (see **Figure 1**). In all experiments but one (Exp. 2-6) those confidence judgments were elicited in an incentive-compatible way (Ducharme and Donnell, 1973; Lebreton et al., 2018a, 2019; Schlag et al., 2015).

**Figure 1.**
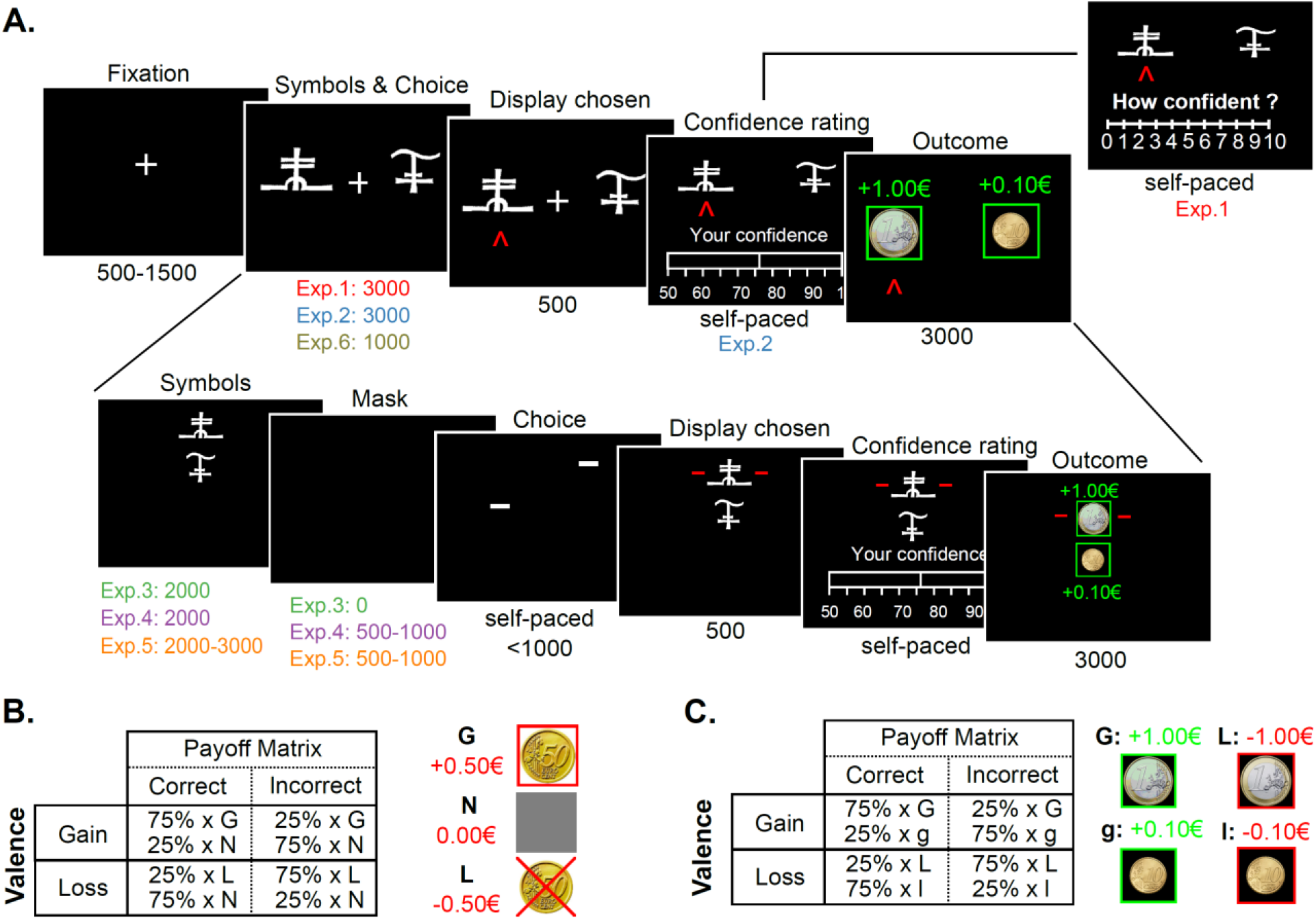
Experimental design. **(A)** Behavioral tasks for Experiments 1-6. Successive screens displayed in one trial are shown from left to right with durations in ms. All tasks are based on the same principle, originally designed for experiments 1-2 (top line): after a fixation cross, participants are presented with a couple of abstract symbols displayed on a computer screen and have to choose between them. They are thereafter asked to report their confidence in their choice on a numerical scale. Outcome associated with the chosen symbol is revealed, sometimes paired with the outcome associated with the unchosen symbol -depending on the condition. For experiments 3-5 (bottom line), options are displayed on a vertical axis. Besides, the response mapping (how the left vs right arrow map to the upper vs lower symbol) is only presented after the symbol display, and the response has to be given within one second of the response mapping screen onset. A short empty screen is used as a mask, between the symbol display and the response mapping for experiment 4-5. Experiment 6 is similar to experiment 2 (top line), except that a shorter duration is allowed from the symbol presentation to the choice Tasks specificities are indicated below each screen. See also **Figure S1** for a complete overview of all 6 experiments. **(B)** Experiment 1 payoff matrix. **(C)** Experiments 2-6 payoff matrix.

Results from experiment 1 and 2 were previously reported in (Lebreton et al., 2019): briefly, we found that participants exhibit the same level of choice accuracy in gain and loss contexts, but are less confident in loss contexts. In addition, they appeared to be slower to execute their choices in loss contexts. Here, in order to evaluate the interdependence between the effects of valence on RT and confidence, we successively designed three additional tasks (**Figure 1A** and **Figure S. 1C-E**). In those tasks, we modified the response setting to blur the effects of valence on RT, with the goal to assess the effects of valence on confidence in the absence of an effect on RT. In a sixth task we imposed a strict time pressure on decisions (**Figure 1A** and **Figure S. 1.F**). All subjects also performed a Transfer task (Lebreton et al., 2019; Palminteri et al., 2015). Data from this additional task is not relevant for our main question of interest and is therefore not analyzed in the present manuscript.

### Learning task - Details

All tasks were implemented using MatlabR2015a^®^ (MathWorks) and the COGENT toolbox (http://www.vislab.ucl.ac.uk/cogent.php). In all experiments, the main learning task was adapted from a probabilistic instrumental learning task used in a previous study (Palminteri et al., 2015). Invited participants were first provided with written instructions, which were reformulated orally if necessary. They were explained that the aim of the task was to maximize their payoff and that gain seeking and loss avoidance were equally important. In each of the three learning sessions, participants repeatedly faced four pairs of cues - taken from Agathodaimon alphabet. The four cue pairs corresponded to four conditions, and were presented 24 times in a pseudo-randomized and unpredictable manner to the subject (intermixed design). Of the four conditions, two corresponded to reward conditions, and two to loss conditions. Within each pair, and depending on the condition, the two cues of a pair were associated with two possible outcomes (1€/0€ for the gain and −1€/0€ for the loss conditions in Exp. 1; 1€/0.1€ for the gain and −1€/-0.1€ for the loss conditions in Exp. 2-6) with reciprocal (but independent) probabilities (75%/25% and 25%/75%) - see (Lebreton et al., 2019) for a detailed rationale.

Experiments 1, 2 and 6 were very similar (**Figure 1A** and **Figure S. 1A-B**): at each trial, participants first viewed a central fixation cross (500-1500ms). Then, the two cues of a pair were presented on each side of this central cross. Note that the side in which a given cue of a pair was presented (left or right of a central fixation cross) was pseudo-randomized, such as a given cue was presented an equal number of times on the left and the right of the screen. Subjects were required to select between the two cues by pressing the left or right arrow on the computer keyboard, within a 3000ms (Exp. 1-2) or 1000ms (Exp. 6) time window. After the choice window, a red pointer appeared below the selected cue for 500ms. Subsequently, participants were asked to indicate how confident they were in their choice. In Experiment 1, confidence ratings were simply given on a rating scale without any additional incentivization. To perform this rating, they could move a cursor –which appeared at a random position- to the left or to the right using the left and right arrows, and validate their final answer with the spacebar. This rating step was self-paced. Finally, an outcome screen displayed the outcome associated with the selected cue, accompanied with the outcome of the unselected cue if the pair was associated with a complete-feedback condition.

In experiment 3, we dissociated the option display and motor response: symbols were first presented on a vertical axis (2s), during this period, participants could choose their preferred symbol, but were uncertain about which button to press to select their preferred symbol. This uncertainty was resolved in the next task phase, in which two horizontal cues indicated which of the left vs right response button could be used to select the top vs bottom symbol (**Figure 1A** and **Figure S.1C**). In addition, we imposed a time limit on the response selection (<1s), to incentivize participants to make their decision during the symbol presentation, and allow only an execution of a choice that was already made during the response mapping screen. In Experiment 4, we added a mask (empty screen 0.5-1s) between the symbol presentation and the response mapping (**Figure 1A** and **Figure S.1D**). This further strengthened the encouragement to make a decision during the symbol presentation to reduce task load, because participants would then only have to retain the information about the selected location (top vs bottom) during the mask period. In Experiment 5, we introduced a jitter (variable time duration; 2-3s) at the symbol presentation screen (**Figure 1A** and **Figure S. 1E**) to further discourage temporal expectations and motor preparedness during the decision period. Finally, Experiment 6 was adapted from Experiment 2, but additionally imposed a strict time pressure on the choice, in an attempt to incentive participants to counteract the slowing down due to the presence of losses (**Figure 1A** and **Figure S. 1E**). In all experiments, response time is defined as the time between the onset of the screen conveying the response mapping (Symbol for Exp. 1-2 & 6; Choice for Exp. 3-5; see **Figure 1A** and **Figure S.1**), and the key press by the participant.

### Matching probability and incentivization

In Experiment 2-6, participant’s reports of confidence were incentivized via a matching probability procedure that is based on the Becker-DeGroot-Marshak (BDM) auction (Becker et al., 1964) Specifically, participants were asked to report as their confidence judgment their estimated probability (p) of having selected the symbol with the higher average value, (i.e. the symbol offering a 75% chance of gain (G75) in the gain conditions, and the symbol offering a 25% chance of loss (L25) in the loss conditions) on a scale between 50% and 100%. A random mechanism, which draws a number (r) in the interval [0.5 1], is then implemented to select whether the subject will be paid an additional bonus of 5 euros as follows: If p ≥ r, the selection of the correct symbol will lead to a bonus payment; if p < r, a lottery will determine whether an additional bonus is won. This lottery offers a payout of 5 euros with probability r and 0 with probability 1-r. This procedure has been shown to incentivize participants to truthfully report their true confidence regardless of risk preferences (Hollard et al., 2016; Karni, 2009). Participants were trained on this lottery mechanism and informed that up to 15 euros could be won and added to their final payment via the MP mechanism applied on one randomly chosen trial at the end of each learning session (3×5 euros). Therefore, the MP mechanism screens were not displayed during the learning sessions.

### Variables

In all experiments, response time is defined as the time between the onset of the screen conveying the response mapping (Symbol for Exp. 1-2 & 6; Choice for Exp. 3-5; see **Figure 1A and Figure S. 1**), and the key press by the participant. Confidence ratings in Exp. 1 were transformed form their original scale (0-10) to a probability scale, (50-100 %), using a simple linear mapping: confidence = (50 + 5 × rating)/100;

### Statistics

All statistical analyses were performed using Matlab R2015a. All reported p-values correspond to twosided tests. T-tests refer to a one sample t-test when comparing experimental data to a reference value (e.g. chance: 0.5), and paired t-tests when comparing experimental data from different conditions. Two-way repeated measures ANOVAs testing for the role of valence, information and their interaction were performed at the individual experiment level.

1-way ANOVAs were used on main effects (e.g. individual averaged accuracy in gains minus losses) to test for the effect of experiments.

Generalized linear mixed-effect (glme) models include a full subject-level random-effects structure (intercepts and slopes for all predictor variables). The models were estimated using Matlab’s fitglme function, which maximize the maximum pseudo-likelihood of observed data under the model (Matlab’s default option). Choice accuracy was modelled using a binomial response function distribution (logistic regression), whereas confidence judgments and response times were modelled using a Normal response function distribution (linear regression).

For instance, the linear mixed-effect models for choice accuracy can be written in Wilkinson-Rogers notation as:

Choice_accuracy ∼ 1 + Val. + Inf. + Val. * Inf. + Fix. + Stim. + Mask. + Sess. + (1 + Val. + Inf. + Val. * Inf. + Fix. + Stim. + Mask. + Sess. |Subject),

With Val: valence; Inf: information; Fix.: fixation duration (only available in Experiments 4-5); Stim.; stimulus display duration (only available in Experiment 5); Mask: Mask duration (only available in Experiments 4-5); Sess: session number.

Note that Val. and Inf. are coded as 0/1, but that the interaction term Val*Inf was computed with Val. and Inf. coded as −1/1 and then rescaled to 0/1.

## Results

First, we evaluated the effects of our manipulation of the display and response settings across the experiments on average levels of choice accuracy and confidence ratings using multiple independent one-way ANOVAs. We found significant effects of the experiments on the average levels of accuracy (F(5,102) = 5.72, P = 1.00×10^-4^, η^2^= 0.21), mostly driven by a drop of accuracy in experiment 6 (see **Table 1** and **Figure S. 2**), but no effects on average levels of confidence ratings (F(5,102) = 1.50, P = 0.1953, η^2^= 0.07; **Table 1**). We also computed, at the session level (participants underwent 3 separate learning sessions per experiment), the correlations between confidence ratings and RT. When averaged at the individual level and tested at the population level (one sample t-test), this measure of the linear relationship between RT and confidence was very significant in all experiments (Exp. 1-6: all Ps < 0.01; **Table 1**). The consistent negative and significant correlations across six experiments indicate that confidence is robustly associated with RT regardless of option-action mapping or time pressure manipulations, suggesting a strong link between instrumental and metacognitive processes. Yet, the correlation between confidence and RT was modulated by our experimental manipulations (effect of experiment: F(5, 102) = 9.91, P < 0.001, η^2^ = 0.32) – post-hoc tests revealed that it was significantly altered by all our experimental manipulations in Exp. 3-6 (**Figure S. 2**).

Next, we analyzed the effects of our experimental manipulation (valence and information) on the observed behavioral variables (choice accuracy, confidence, RT), using repeated measures ANOVAs in each individual study (**Figure 2**; **Table 2**). The parallel analyses of choice accuracy and confidence ratings replicated the results reported in (Fontanesi et al., 2019; Lebreton et al., 2019; Palminteri et al., 2015). Indeed, participants were more accurate in complete information contexts in five of six experiments (**Table 2**; main effect of information on accuracy, Exp. 1-5: Ps < 0.05; Exp. 6: P = 0.1570)). The effects of information on accuracy were actually not significantly different across our different experiments (**Figure S. 3**; effect of experiment: F(102,5) = 0.52; P = 0.7289 η^2^ = 0.03). On the other hand, participants learned equally well in gain and loss contexts, as they exhibited similar levels of accuracy in gains and loss contexts in all experiments (**Table 2**; main effect of valence on accuracy, Exp. 1-6: all Ps > 0.3; **Figure S. 3;** effect of experiment: F(5, 102) = 0.35, P = 0.884, η^2^ = 0.02).

**Figure 2.**
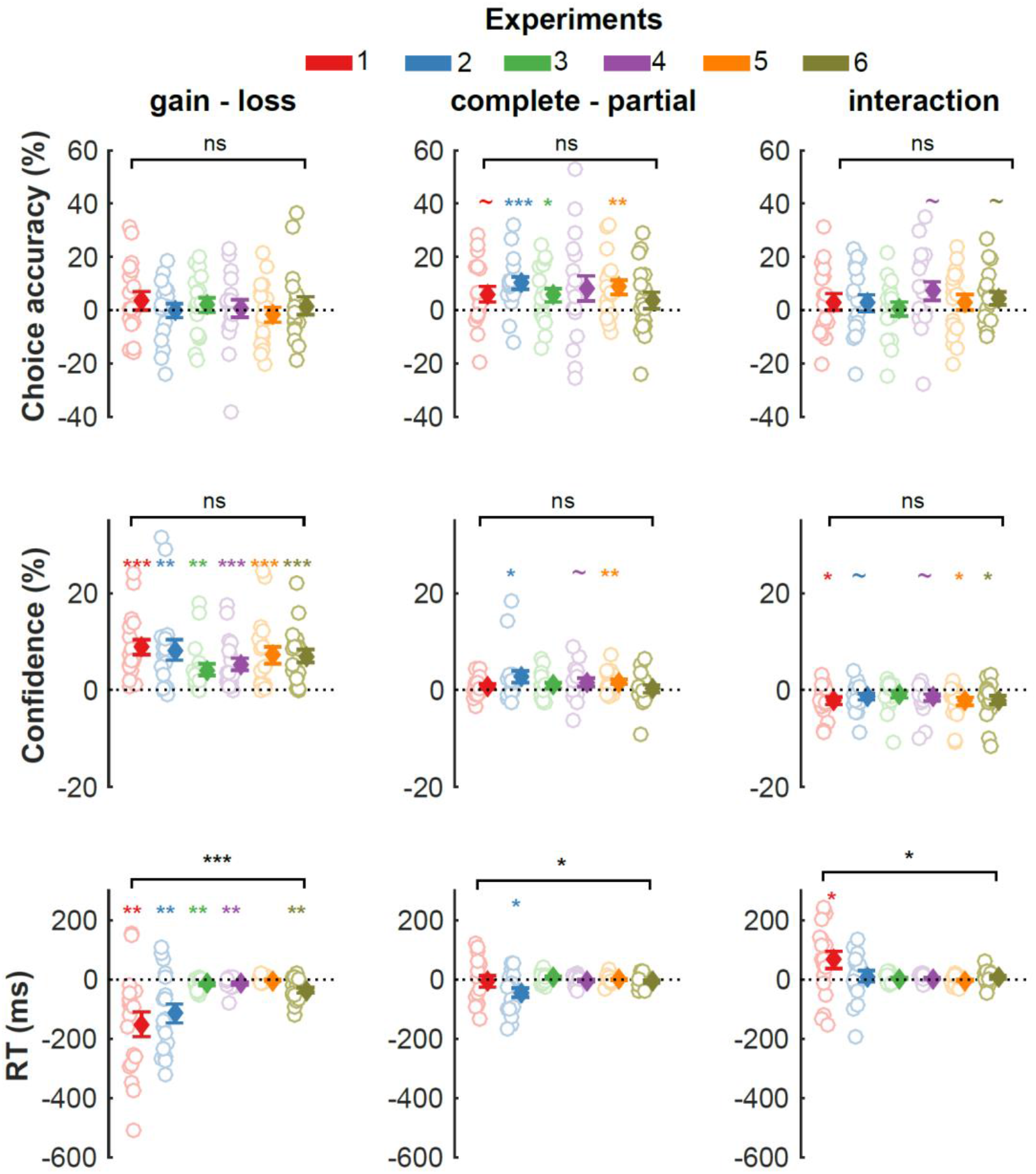
Behavioral results. Effects of the main manipulations (left: valence; middle: information; right: interaction) on relevant measures of choice-relevant behavior (top: performance; middle: confidence; bottom: response times). Analyses are independently performed in the six different experiments using repeated-measures ANOVAs. Empty dots with colored edges represent individual data points across different experiments; filled diamonds and error-bars represent sample mean ± SEM. The horizontal bar indicates a one-way ANOVA testing the effect of experiment on each manipulation (see supplementary materials, **Figure S. 3** for details). ∼ P<0.1; * P<0.05; ** P<0.01; *** P<0.001

**Table 2.** Repeated mgeasures ANOVA results reported separately for choice-relevant behavioral measures. val: valence; inf: information; ∼ P<0.1; * P<0.05; ** P<0.01; *** P<0.001

|  |  |  | Exp.1 | Exp. 2 | Exp. 3 | Exp. 4 | Exp. 5 | Exp. 6 |
| --- | --- | --- | --- | --- | --- | --- | --- | --- |
| Performance | val. | F(1,17), [$\eta^2$]<br>(P-val.) | 1.04, [0.01]<br>(0.323) | 0.00, [0.00]<br>(0.971) | 0.40, [0.00]<br>(0.538) | 0.01, [0.00]<br>(0.912) | 0.33, [0.00]<br>(0.571) | 0.37, [0.04]<br>(0.553) |
| | inf. | F(1,17), [$\eta^2$]<br>(P-val.) | 4.28, [0.04]<br>(0.054)~ | 18.64, [0.15]<br>(0.001)*** | 5.56, [0.04]<br>(0.031)* | 3.26, [0.06]<br>(0.089)~ | 10.17, [0.07]<br>(0.005)** | 2.19, [0.02]<br>(0.157) |
| | val×inf | F(1,17), [$\eta^2$]<br>(P-val.) | 1.06, [0.01]<br>(0.319) | 0.77, [0.01]<br>(0.393) | 0.06, [0.00]<br>(0.816) | 4.36, [0.04]<br>(0.052)~ | 1.04, [0.01]<br>(0.326) | 3.57, [0.02]<br>(0.075)~ |
| Confidence | val. | F(1,17), [$\eta^2$]<br>(P-val.) | 33.11, [0.27]<br>($<0.001$)*** | 15.43, [0.19]<br>(0.001)** | 12.18, [0.03]<br>(0.003)** | 19.14, [0.07]<br>($<0.001$)*** | 16.71, [0.15]<br>($<0.001$)*** | 26.71, [0.12]<br>($<0.001$)*** |
| | inf. | F(1,17), [$\eta^2$]<br>(P-val.) | 2.00, [0.00]<br>(0.175) | 4.92, [0.02]<br>(0.040)* | 2.28, [0.02]<br>(0.149) | 3.21, [0.01]<br>(0.091)~ | 11.07, [0.01]<br>(0.004)** | 0.11, [0.00]<br>(0.743) |
| | val×inf | F(1,17), [$\eta^2$]<br>(P-val.) | 7.58, [0.02]<br>(0.014)* | 4.25, [0.01]<br>(0.055)~ | 1.61, [0.01]<br>(0.222) | 4.46, [0.01]<br>(0.050)~ | 7.87, [0.02]<br>(0.012)* | 5.16, [0.01]<br>(0.036)* |
| RT | val. | F(1,17), [$\eta^2$]<br>(P-val.) | 13.25, [0.03]<br>(0.002)** | 13.15, [0.08]<br>(0.002)** | 12.47, [0.01]<br>(0.003)** | 11.23, [0.01]<br>(0.004)** | 1.97, [0.00]<br>(0.178) | 15.56, [0.02]<br>(0.001)** |
| | inf. | F(1,17), [$\eta^2$]<br>(P-val.) | 0.12, [0.00]<br>(0.733) | 7.64, [0.01]<br>(0.013)* | 1.82, [0.00]<br>(0.195) | 0.31, [0.00]<br>(0.586) | 0.09, [0.00]<br>(0.766) | 3.60, [0.00]<br>(0.074)~ |
| | val×inf | F(1,17), [$\eta^2$]<br>(P-val.) | 4.94, [0.01]<br>(0.040)* | 0.36, [0.00]<br>(0.558) | 1.32, [0.00]<br>(0.266) | 2.32, [0.00]<br>(0.146) | 0.70, [0.00]<br>(0.414) | 2.02, [0.00]<br>(0.173) |

Despite similar performances in gain and loss contexts, and despite our attempt to cancel the PIT with our manipulations of the option-action mapping and time pressure, participants were slower in loss contexts in experiments 1-4 & 6 (**Table 2**; main effect of valence on RT: all Ps < 0.01). These results not only replicate the results reported in (Fontanesi et al., 2019), but also assert the robustness of the PIT to the manipulation of response setups in human instrumental learning. Still, our experimental manipulations significantly reduced the PIT in Exp. 3-5 (**Figure S. 3;** effect of experiment: F(5, 102) = 7.98, P < 0.001, η^2^ = 0.28).

Importantly, despite similar performance in gain and loss contexts, participants were less confident in loss contexts (**Table 2**; main effect of valence on confidence, Exp. 1-6: all Ps < 0.01), with very similar effect sizes across all experiments (**Figure S. 3**; F(5, 102) = 1.26, P = 0.289, η^2^ = 0.06). These effects were mitigated when more information was available (**Table 2**; interaction valence × information on confidence: all Ps < 0.05). These results not only replicate those reported in Lebreton et al., 2019, but also assert the robustness of the PMT.

Overall, the analyses of the data collected in six different versions of our experiment (N = 108) clearly underline the remarkable robustness of the effects of outcome valence on both confidence and RT. Only one experimental condition succeeded in cancelling the PIT (Experiment 5). Note that in this experiment, we still observed the PMT as evidenced by a significant main effect of valence on confidence (**Table 2**; F(1,17) = 16.71, P < 0.001, η^2^= 0.15), but not on RT (F(1,17) = 1.97, P=0.178, η^2^= 0.001). This suggests that the effects of outcome valence on confidence and RT are – partly – dissociable. In other words, we can observe a lower confidence in loss contexts, even when RTs are indistinguishable from gain contexts.

In order to give a comprehensive overview of the relationship between accuracy, confidence and RT, and to quantify the effects of the different available predictors on these behavioral measures, we also ran generalized linear mixed-effect regressions. Independent variables included not only valence, information and their interaction, but also the different available timings (e.g. duration of the stimulus or mask display) and a linear trend accounting for the session effects (see methods for details). These sensitive trial-by-trial analyses replicated the main ANOVA results reported above regarding the effects of valence and information on performance, confidence and RT (**Figure 3**; **Tables 4–6**). They also confirmed that, in experiment 5, no effect of valence can be detected on RT and performance (P = 0.349 and P = 0.620) while a robust effect is observed on confidence (P = 0.002).

**Figure 3.**
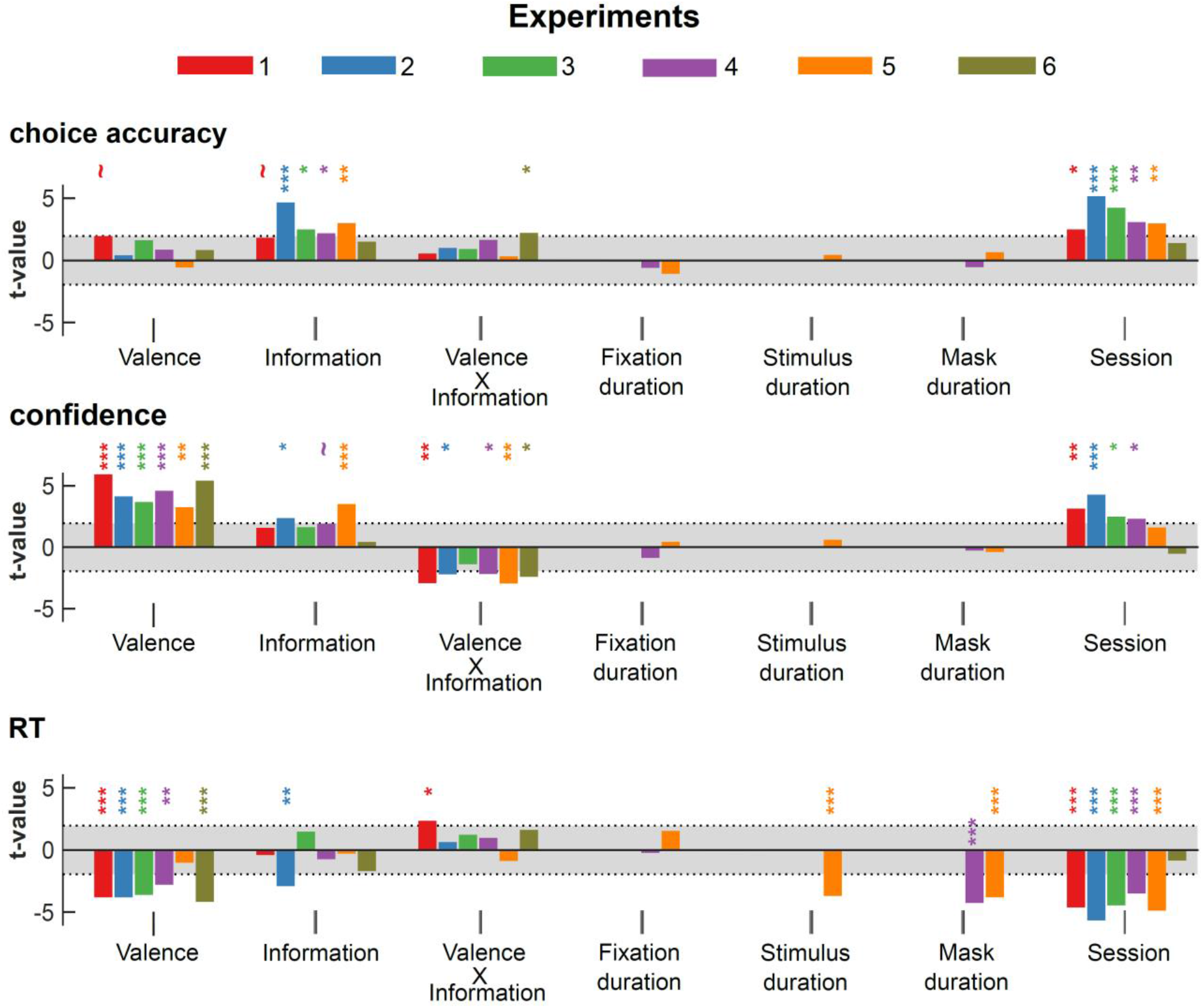
Generalized linear mixed-effects models. Estimated standardized regression coefficients (t-values) from generalized linear mixed-effects (GLME) models, fitted in the different experiments. Top: logistic GLME with performance as the dependent variable. Middle: linear GLME with confidence as the dependent variable. Bottom: linear GLME with RT as the dependent variable; Shaded area represent area where coefficients are not significantly different from 0 (abs(t-value) < 1.95; p>0.05). ∼ P<0.1; * P<0.05; ** P<0.01; *** P<0.001

**Table 3.** Estimated coefficients from inter-individual robust regressions. For each individual, we estimated the net effect of valence on RT and confidence, by computing the averaged difference of these behavioral measures in the gain versus loss contexts. For analyses restricted to a single experiment, we used robust regressions to decrease the vulnerability of our estimates in the relatively small samples (n=18). For the combined analysis (n=108), simple and robust regressions gave similar results, and we only report here the results of the simple regression. β: estimated regression coefficient. SE: estimated standard error of the regression coefficient. ∼ P<0.1; * P<0.05; ** P<0.01; *** P<0.001

|  |  | Exp. 1 | Exp. 2 | Exp. 3 | Exp. 4 | Exp. 5 | Exp. 6 | All |
| --- | --- | --- | --- | --- | --- | --- | --- | --- |
| **Intercept** | $\beta \pm \text{SE}$ | $8.02 \pm 2.15$ | $2.94 \pm 1.29$ | $2.16 \pm 0.95$ | $3.54 \pm 1.37$ | $6.76 \pm 1.81$ | $3.59 \pm 1.95$ | $5.62 \pm 0.69$ |
|  | t-val | 3.72 | 2.27 | 2.27 | 2.58 | 3.73 | 2.41 | 8.18 |
|  | (P-val) | (0.002)** | (0.037)* | (0.038)* | (0.020)* | (0.002)** | (0.028)* | (<0.001)*** |
| **Slope** | $\beta \pm \text{SE}$ | $-0.003 \pm 0.009$ | $-0.03 \pm 0.01$ | $-0.02 \pm 0.04$ | $-0.06 \pm 0.06$ | $0.17 \pm 0.18$ | $-0.10 \pm 0.03$ | $-0.02 \pm 0.01$ |
|  | t-val | -0.27 | -3.55 | -0.46 | -0.97 | 0.81 | -3.35 | -3.75 |
|  | (P-val) | (0.793) | (0.003)** | (0.662) | (0.35) | (0.368) | (0.004)** | (<0.001)*** |

**Table 4.** Estimated coefficients from generalized linear mixed-effect models on performance across experiments. β: estimated regression coefficients for fixed effects. SE: estimated standard error of the regression coefficients. Val: valence; Inf: information; Fix.: fixation duration; Stim.; stimulus display duration; Sess: session number. ∼ P<0.1; * P<0.05; ** P<0.01; *** P<0.001

|  |  | Exp. 1 | Exp. 2 | Exp. 3 | Exp. 4 | Exp. 5 | Exp. 6 |
| --- | --- | --- | --- | --- | --- | --- | --- |
| **Val.** | $\beta \pm \text{SE}$ | $0.40 \pm 0.21$ | $0.08 \pm 0.18$ | $0.16 \pm 0.27$ | $0.15 \pm 0.19$ | $-0.08 \pm 0.17$ | $0.12 \pm 0.15$ |
|  | t-val | 1.86 | 0.32 | 0.58 | 0.78 | -0.50 | 0.76 |
|  | (P-val) | (0.063)~ | (0.748) | (0.561) | (0.43) | (0.620) | (0.449) |
| **Inf.** | $\beta \pm \text{SE}$ | $0.31 \pm 0.18$ | $0.72 \pm 0.16$ | $0.52 \pm 0.22$ | $0.63 \pm 0.30$ | $0.51 \pm 0.18$ | $0.22 \pm 0.16$ |
|  | t-val | 1.74 | 4.59 | 2.40 | 2.10 | 2.92 | 1.41 |
|  | (P-val) | (0.081)~ | (<0.001)*** | (0.016)* | (0.036)* | (0.004)** | (0.158) |
| **Val x Inf** | $\beta \pm \text{SE}$ | $0.10 \pm 0.20$ | $0.20 \pm 0.21$ | $0.16 \pm 0.19$ | $0.36 \pm 0.23$ | $0.04 \pm 0.18$ | $0.25 \pm 0.12$ |
|  | t-val | 0.47 | 0.92 | 0.83 | 1.56 | 0.23 | 2.13 |
|  | (P-val) | (0.638) | (0.356) | (0.405) | (0.118) | (0.814) | (0.034)* |
| **Fix (s)** | $\beta \pm \text{SE}$ | | | | $0.18 \pm 0.35$ | $-0.28 \pm 0.28$ | |
|  | t-val | - | - | - | -0.51 | -0.99 | - |
|  | (P-val) |  |  |  | (0.611) | (0.322) |  |
| **Stim (s)** | $\beta \pm \text{SE}$ | | | | | $0.03 \pm 0.07$ | |
|  | t-val | - | - | - | - | 0.37 | - |
|  | (P-val) |  |  |  |  | (0.713) |  |
| **Mask (s)** | $\beta \pm \text{SE}$ | | | | $0.14 \pm 0.29$ | $0.17 \pm 0.28$ | |
|  | t-val | - | - | - | -0.47 | 0.57 | - |
|  | (P-val) |  |  |  | (0.637) | (0.567) |  |
| **Sess.** | $\beta \pm \text{SE}$ | $0.34 \pm 0.14$ | $0.78 \pm 0.15$ | $0.58 \pm 0.14$ | $0.46 \pm 0.15$ | $0.30 \pm 0.10$ | $0.09 \pm 0.07$ |
|  | t-val | 2.40 | 5.08 | 4.17 | 3.00 | 2.89 | 1.32 |
|  | (P-val) | (0.016)* | (<0.001)*** | (<0.001)*** | (0.003)** | (0.004)** | (0.186) |

**Table 5.**
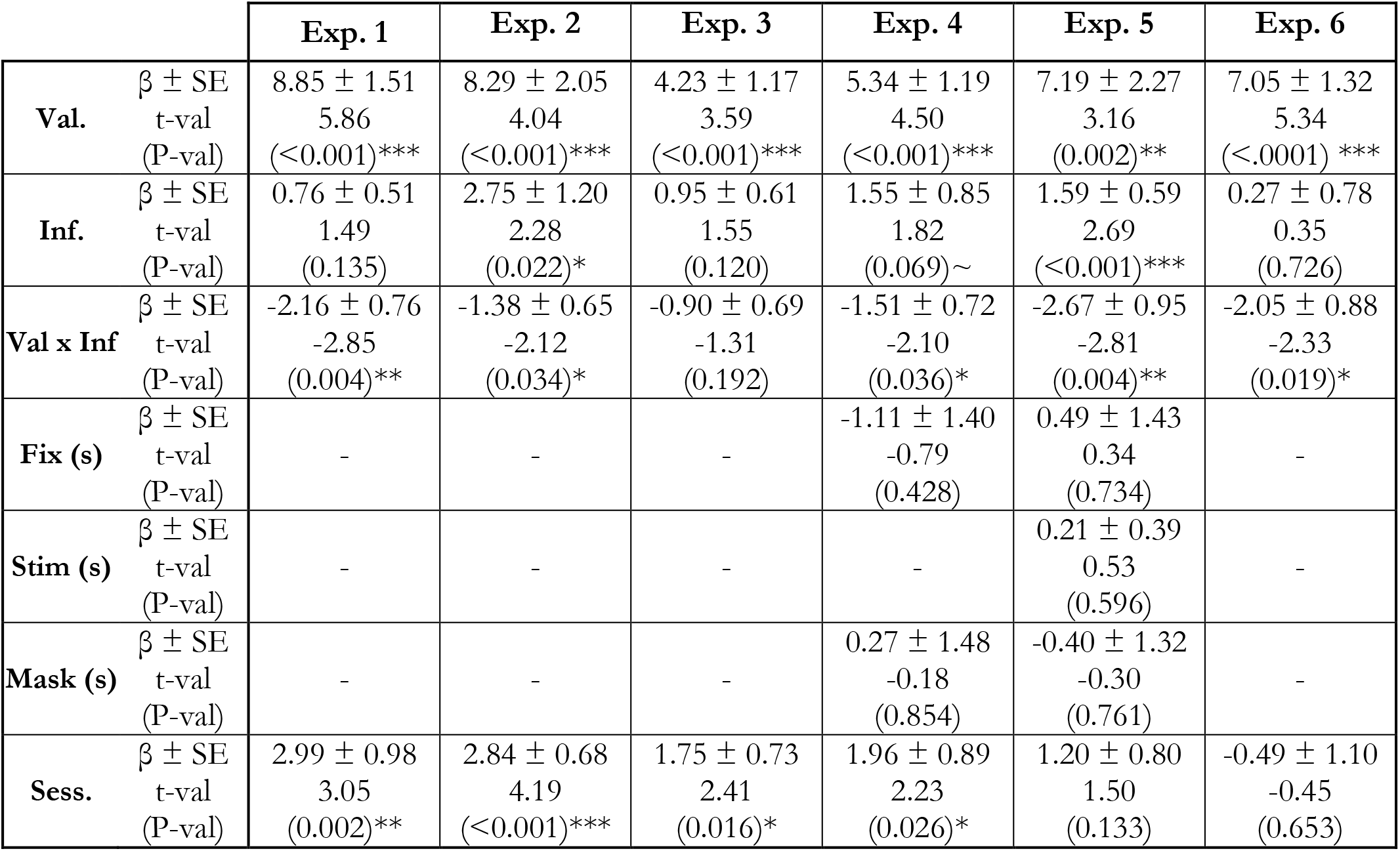
Estimated coefficients from generalized linear mixed-effect models on confidence across experiments. β: estimated regression coefficients for fixed effects. SE: estimated standard error of the regression coefficients. Val: valence; Inf: information; Fix.: fixation duration; Stim.; stimulus display duration; Sess: session number. ∼ P<0.1; * P<0.05; ** P<0.01; *** P<0.001

|  |  | **Exp. 1** | **Exp. 2** | **Exp. 3** | **Exp. 4** | **Exp. 5** | **Exp. 6** |
| --- | --- | --- | --- | --- | --- | --- | --- |
| **Val.** | $\beta \pm \text{SE}$ | $8.85 \pm 1.51$ | $8.29 \pm 2.05$ | $4.23 \pm 1.17$ | $5.34 \pm 1.19$ | $7.19 \pm 2.27$ | $7.05 \pm 1.32$ |
|  | t-val | 5.86 | 4.04 | 3.59 | 4.50 | 3.16 | 5.34 |
|  | (P-val) | (<0.001)*** | (<0.001)*** | (<0.001)*** | (<0.001)*** | (0.002)** | (<.0001) *** |
| **Inf.** | $\beta \pm \text{SE}$ | $0.76 \pm 0.51$ | $2.75 \pm 1.20$ | $0.95 \pm 0.61$ | $1.55 \pm 0.85$ | $1.59 \pm 0.59$ | $0.27 \pm 0.78$ |
|  | t-val | 1.49 | 2.28 | 1.55 | 1.82 | 2.69 | 0.35 |
|  | (P-val) | (0.135) | (0.022)* | (0.120) | (0.069)~ | (<0.001)*** | (0.726) |
| **Val x Inf** | $\beta \pm \text{SE}$ | $-2.16 \pm 0.76$ | $-1.38 \pm 0.65$ | $-0.90 \pm 0.69$ | $-1.51 \pm 0.72$ | $-2.67 \pm 0.95$ | $-2.05 \pm 0.88$ |
|  | t-val | -2.85 | -2.12 | -1.31 | -2.10 | -2.81 | -2.33 |
|  | (P-val) | (0.004)** | (0.034)* | (0.192) | (0.036)* | (0.004)** | (0.019)* |
| **Fix (s)** | $\beta \pm \text{SE}$ | | | | $-1.11 \pm 1.40$ | $0.49 \pm 1.43$ | |
|  | t-val | - | - | - | -0.79 | 0.34 | - |
|  | (P-val) |  |  |  | (0.428) | (0.734) |  |
| **Stim (s)** | $\beta \pm \text{SE}$ | | | | | $0.21 \pm 0.39$ | |
|  | t-val | - | - | - | - | 0.53 | - |
|  | (P-val) |  |  |  |  | (0.596) |  |
| **Mask (s)** | $\beta \pm \text{SE}$ | | | | $0.27 \pm 1.48$ | $-0.40 \pm 1.32$ | |
|  | t-val | - | - | - | -0.18 | -0.30 | - |
|  | (P-val) |  |  |  | (0.854) | (0.761) |  |
| **Sess.** | $\beta \pm \text{SE}$ | $2.99 \pm 0.98$ | $2.84 \pm 0.68$ | $1.75 \pm 0.73$ | $1.96 \pm 0.89$ | $1.20 \pm 0.80$ | $-0.49 \pm 1.10$ |
|  | t-val | 3.05 | 4.19 | 2.41 | 2.23 | 1.50 | -0.45 |
|  | (P-val) | (0.002)** | (<0.001)*** | (0.016)* | (0.026)* | (0.133) | (0.653) |

**Table 6.** Estimated coefficients from generalized linear mixed-effect models on response times across experiments. β: estimated regression coefficients for fixed effects. SE: estimated standard error of the regression coefficients. Val: valence; Inf: information; Fix.: fixation duration; Stim.; stimulus display duration; Sess: session number. ∼ P<0.1; * P<0.05; ** P<0.01; *** P<0.001

|  |  | Exp. 1 | Exp. 2 | Exp. 3 | Exp. 4 | Exp. 5 | Exp. 6 |
| --- | --- | --- | --- | --- | --- | --- | --- |
| **Val.** | $\beta \pm SE$ | $-151.12 \pm 40.37$ | $-115.63 \pm 30.96$ | $-15.31 \pm 4.33$ | $-13.49 \pm 4.97$ | $-3.23 \pm 3.44$ | $-35.19 \pm 8.60$ |
|  | t-val | -3.74 | -3.73 | -3.53 | -2.71 | -0.94 | -4.10 |
|  | (P-val) | (<0.001)*** | (<0.001)*** | (<0.001)*** | (0.007)** | (0.349) | (<0.001)*** |
| **Inf.** | $\beta \pm SE$ | $-6.57 \pm 19.58$ | $-44.37 \pm 15.75$ | $5.81 \pm 4.13$ | $-2.81 \pm 4.28$ | $0.80 \pm 3.78$ | $-8.23 \pm 5.11$ |
|  | t-val | -0.34 | -2.82 | 1.41 | -0.65 | -0.21 | -1.61 |
|  | (P-val) | (0.737) | (0.005)** | (0.160) | (0.513) | (0.832) | (0.107) |
| **Val x Inf** | $\beta \pm SE$ | $65.58 \pm 28.77$ | $10.59 \pm 18.88$ | $3.75 \pm 3.25$ | $3.67 \pm 4.19$ | $-3.04 \pm 3.85$ | $8.35 \pm 5.43$ |
|  | t-val | 2.28 | 0.56 | 1.15 | 0.88 | -0.79 | 1.54 |
|  | (P-val) | (0.023)* | (0.575) | (0.249) | (0.381) | (0.430) | (0.124) |
| **Fix (s)** | $\beta \pm SE$ | | | | $-2.37 \pm 16.13$ | $18.56 \pm 12.77$ | |
|  | t-val | - | - | - | -0.15 | 1.45 | - |
|  | (P-val) |  |  |  | (0.883) | (0.146) |  |
| **Stim (s)** | $\beta \pm SE$ | | | | | $-13.12 \pm 4.00$ | |
|  | t-val | - | - | - | - | -3.28 | - |
|  | (P-val) |  |  |  |  | (<0.001)*** |  |
| **Mask(s)** | $\beta \pm SE$ | | | | $-68.34 \pm 16.35$ | $-54.20 \pm 14.5$ | |
|  | t-val | - | - | - | -4.18 | -3.73 | - |
|  | (P-val) |  |  |  | (<0.001)*** | (<0.001)*** |  |
| **Sess.** | $\beta \pm SE$ | $-152.43 \pm 33.63$ | $-146.28 \pm 26.13$ | $-26.93 \pm 6.14$ | $-32.55 \pm 9.51$ | $-27.57 \pm 5.64$ | $-6.39 \pm 8.34$ |
|  | t-val | -4.53 | -5.60 | -4.38 | -3.42 | -4.79 | -0.77 |
|  | (P-val) | (<0.001)*** | (<0.001)*** | (<0.001)*** | (<0.001)*** | (<0.001)*** | (0.443) |

Because the PIT – i.e. the slowing down of RTs in loss compared to gain contexts – was extremely robust to our experimental manipulations aiming at cancelling it, the ANOVA and regressions above provide only limited evidence on whether valence-induced decreasing on confidence can be observed in the absence of the valence-induced slowing of RT. In the following paragraphs, we therefore used a different analytical strategy leveraging inter-individual differences to test this hypothesis. We assessed the link between individual slowing down (RT in gain – loss) and individual decreases in confidence (confidence in gain-loss) in our full sample and in each individual study using robust linear regressions (see methods for details). In those regressions, the coefficients for the intercept and slope quantify two different but equally important signals: First, the y-intercept represents a theoretical individual who exhibits no effect of valence on RT (RT in gain – loss = 0, Figure 4A): an intercept significantly different from 0 therefore indicates that a significant effect of valence on confidence can be observed in the absence of an effect on RT. Second, the slope quantifies how the effect of valence on confidence linearly depends on the valence-induced slowing of RT. Both at the population level (i.e., combining data from all six experiments) and in each individual study, the intercepts of those regressions were estimated to be significantly positive (all Ps < 0.05; Figure 4A-B; Table 4). This indicates that valence-induced changes on confidence are detectable when valence induced-changes on RT are absent. Note that at the population level, the slope of the regression was also significantly negative (β = −0.02 ± 0.01, t(106) = −3.75, P < 0.001), indicating that, compared with the gain context, the more participants were slowed down by the loss context, the less confident they were in their response. Therefore, PMT and PIT are only partially dissociable.

**Figure 4.**
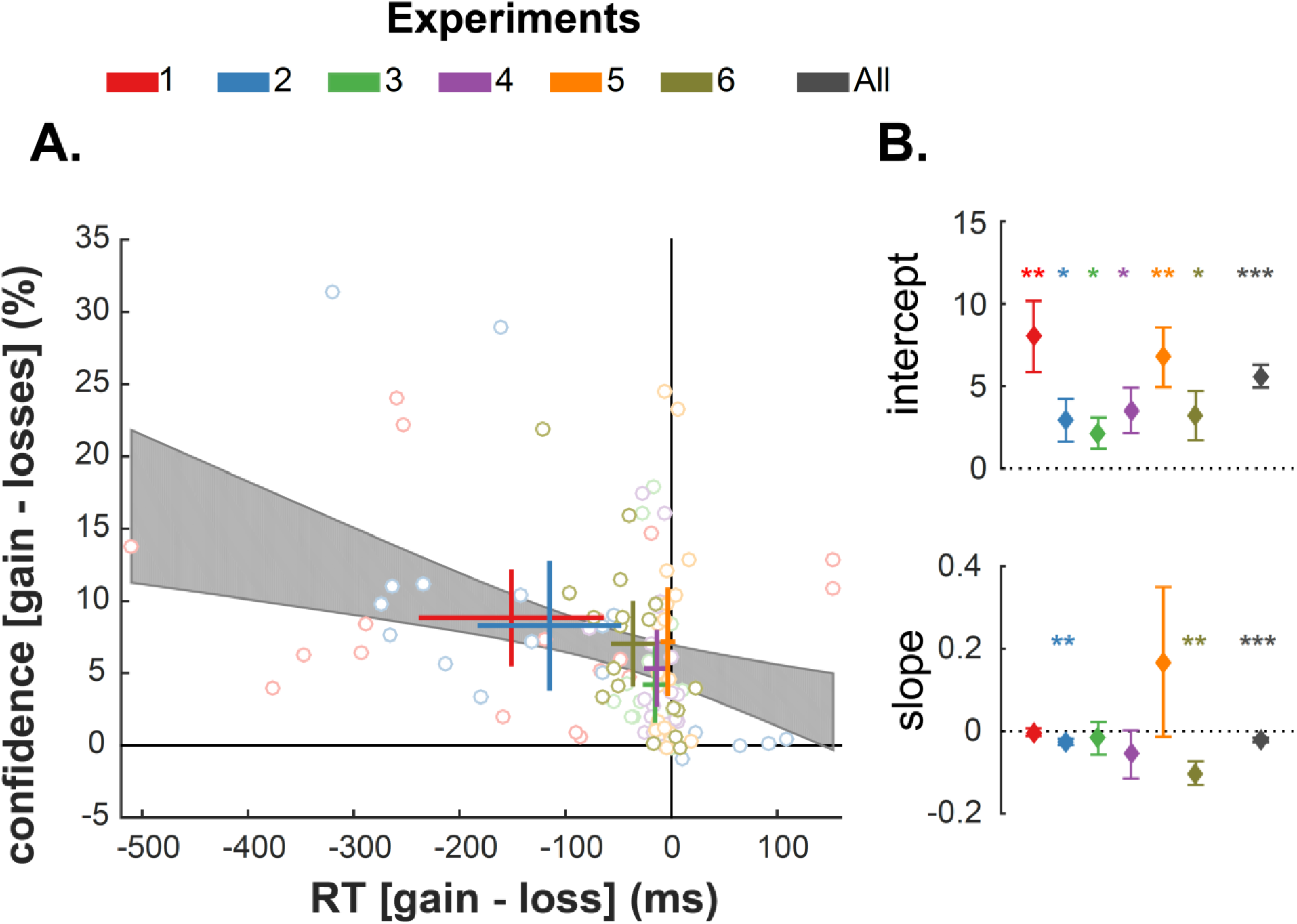
Assessing the link between the effects of valence on confidence and response times. **(A)** Interindividual correlations between the effects of valence on confidence (Y-axis) and response times (X-axis) across experiments. Dots represent data points from individual participants. Thick lines represent the mean ± 95%CI of the effects of valence on confidence (vertical lines) and response times (horizontal lines). Experiments are indicated by the dot edge and line color. The black shaded area represents the 95%CI of the inter-individual linear regression. Note that potential outliers did not bias the regression, given that simple and robust regressions gave very similar results. **(B)** Results from inter-individual regressions of the valence-induced RT slowing on the valence-induced confidence difference across different experiments. Top: estimated intercepts of the regressions. Bottom: estimated slopes of the regressions. Diamonds and error-bars represent the estimated regression coefficients (β) and their standard error. * P<0.05; ** P<0.01; *** P<0.001

## Discussion

The present work investigated the relationship between valence-induced biases affecting two different behavioral outputs: response time and confidence. In simple probabilistic reinforcement-learning tasks, learning to avoid punishment increased participants’ response time (RT) and decreased their confidence in their choices, without affecting their actual performance (Fontanesi et al., 2019; Lebreton et al., 2019). The valence-induced bias on RT is currently interpreted as a manifestation of the Pavlovian-Instrumental Transfer phenomenon (Boureau and Dayan, 2011; Guitart-Masip et al., 2012). The valence-induced decrease in confidence – akin to a Pavlovian-Metacognition Transfer – has been described as a value-to-confidence contamination, potentially generated by a mechanisms of affect-as-information (Lebreton et al., 2018; Schwarz and Clore, 1983).

One of the motivations behind the present study was to rule out a potential alternative explanation of the observed decrease in confidence: participants could derive confidence estimates by monitoring changes in their own response times. Indeed, because it has been suggested that humans can infer confidence levels from observing their RT (Desender et al., 2017; Kiani et al., 2014), the valence-induced bias on confidence could be spuriously driven by a Pavlovian-Instrumental Transfer mechanism operating at the level of motor initiation (Boureau and Dayan, 2011; Guitart-Masip et al., 2012). As such, valence-induced confidence biases would then merely reflect a secondary effect of valence mediated by response time slowing, and not a primary meta-cognitive bias. Crucially, this possibility is not ruled out by previous studies, where effects of affective states on confidence judgments in perceptual or cognitive tasks typically lacked control over RT (Giardini et al., 2008; Koellinger and Treffers, 2015; Massoni, 2014, but see Lebreton et al., 2018). We address this issue in the current set of experiments by dissociating decisions from motor mapping, thereby partially removing the association between RT and confidence.

We analyzed six datasets composed of two published datasets (Exp. 1-2) and four new experiments (Exp. 3-6). Over those six experimental datasets, the first noticeable result is that we systematically replicated previous instrumental learning results using the same paradigm with very consistent effect sizes (Palminteri et al., 2015, 2016): participants learn equally well to seek reward and avoid punishment, and learning performance benefits from complete information (i.e. feedback about the counterfactual outcome). The reliability of the results extended beyond choice behavior as confidence and RT were, respectively, lower and slower in punishment contexts compared to reward contexts, as previously reported (Fontanesi et al. 2019; Lebreton et al., 2019), thus confirming the robustness of the valence bias.

The second important result is that the slowing down of RTs in loss contexts is extremely resilient, as it was still observed when the mapping between motor response and option selection was dissociated by our experimental design (Exp. 3-4) and when significant time pressure was applied on the decision (Exp. 6) – albeit with significantly lower effect sizes. This result speaks to the strength and the pervasiveness of the Pavlovian-Instrumental Transfer mechanism operating at the motor level (Boureau and Dayan, 2011; Guitart-Masip et al., 2012).

Third, and importantly, we still observed a significant valence effect on confidence when the valence effects on RT were dramatically reduced (Exp. 3, 4 and 6) or absent (Exp. 5), indicating that the lower confidence observed in the loss-avoidance context (PMT) is – at least partly– dissociable from the purely motor components of PIT. This was confirmed by additional evidence from inter-individual difference analyses, showing that in all six experiments, a theoretical subject exhibiting no valence-induced bias in RT would still exhibit a valence-induced bias in confidence. Altogether, these results suggests that it is unlikely that the valence-induced bias on confidence reported in human reinforcement-learning (Lebreton et al., 2019) is a mere consequence of a response time slowing caused by a Pavlovian-Instrumental Transfer. Our results are also consistent with recent findings (Dotan et al., 2018) challenging the notion that humans infer confidence levels purely from observing their own response times, and suggesting that decision reaction times are a consequence rather than a cause of the feeling of confidence (Desender et al., 2017; Kiani et al., 2014). It is worth noting that in most studies, decision-time (i.e. when participants reach a decision) and response times (when participants indicate their choice) are not experimentally dissociated and often conflated in the same measure. Here we delayed the mapping between decisions (in the option space) and action selection (motor space), which resulted in an effective control over response times. Future studies will investigate whether participants can keep track of an internal measure of decision time, which could influence confidence.

In our data, we also observed that confidence ratings and RT are robustly associated regardless of time pressure manipulation. The negative correlation between confidence and RT was consistently found in over six experiments. This coupling is consistent with predictions from most of sequential-sampling model (van den Berg et al., 2016; De Martino et al., 2013; Navajas et al., 2016; Pleskac and Busemeyer, 2007; Ratcliff and Starns, 2009, 2013; Yu et al., 2015), which posit that confidence and RT jointly emerge from a single mechanism of evidence accumulation. Importantly, we still observed robust correlation between confidence and motor RTs when we dissociated action selection from the option evaluation. Therefore, the motor execution of a decision might be more important than previously thought in sequential-sampling models of confidence, which mostly focus on decision times.

The replicability and robustness of the Pavlovian-Metacognition Transfer implies that the manipulations of valence could prove useful to dissociate fundamental components of decisionmaking and metacognitive judgment, such as objective uncertainty and subjective confidence (Bang and Fleming, 2018). The dissociation between objective uncertainty and subjective confidence is anticipated by post-decisional and second-order models of confidence (Fleming and Daw, 2017; Pleskac and Busemeyer, 2007), which postulate that confidence is formed after the decision and thereby might be influenced by other internal or external variables (Moran et al., 2015; Navajas et al., 2016; Yu et al., 2015). It is worth noting that our results do not rule out the possibility that RT is used to guide metacognitive judgment of confidence before and after the decision. Actually, the fact that participants who exhibit the strongest PIT are also the ones that exhibit the strongest PMT (significant negative slope(s) in **Figure 4A-B** and **Table 3**) indicates that their reaction times and confidence are linked. Observing one’s RTs could therefore be one of the factors that influences confidence after the decision was made, as posited in second-order models.

In a previous study (Fontanesi et al., 2019), we analyzed the effects of valence on RT, on a different dataset collected with a similar experimental design – although omitting confidence judgments. There, using an approach combining reinforcement-learning and decision-diffusion modelling, we reported that valence influences two critical parameters of the response time model: the non-decision-time - which typically represents perceptual and motor processes – and the decision threshold – which indexes response cautiousness. We speculate that this distinction is relevant to interpret the results of the present report. We propose that the portion of the valence-induced response time slowing that we were able to cancel through response-mapping manipulation could be linked to the non-decision-time modulation; on the other hand the residual irreducible valence-induced response time slowing could be linked to the increased response cautiousness. Yet, given the disruption of the response mapping present in most experiments in the current study, the combined reinforcement-learning and decision-diffusion modelling approach cannot be applied to the present data to test this hypothesis. Further experiments are therefore needed to refine the computational description of valence-induced biases in reinforcement-learning, and their consequences on performance, confidence and response times.

