## Supplementary Material for "Robust Pavlovian-to-Instrumental and Pavlovian-to-Metacognitive Transfers in human reinforcement learning"

### Supplementary Materials

| | | Gain - Loss | Partial - Complete | Interaction |
|---|---|---|---|---|
| Choice Accuracy | F(5, 102), [$\eta^2$] (p-val) | 0.35, [0.02] (0.884) | 0.52, [0.03] (0.729) | 0.56, [0.03] (0.725) |
| Confidence | F(5, 102), [$\eta^2$] (p-val) | 1.26, [0.06] (0.289) | 1.19, [0.06] (0.319) | 0.51, [0.02] (0.771) |
| RT | F(5, 102), [$\eta^2$] (p-val) | 7.98, [0.28] (<0.001) *** | 2.81, [0.12] (0.021)* | 2.98, [0.13] (0.015)* |

**Table S1.** One-way ANOVA results for the combined data from six experiments. The role of experimental manipulations on the effect of outcome valence on Accuracy, Confidence and RT.
~ P<0.1; * P<0.05; ** P<0.01; *** P<0.001

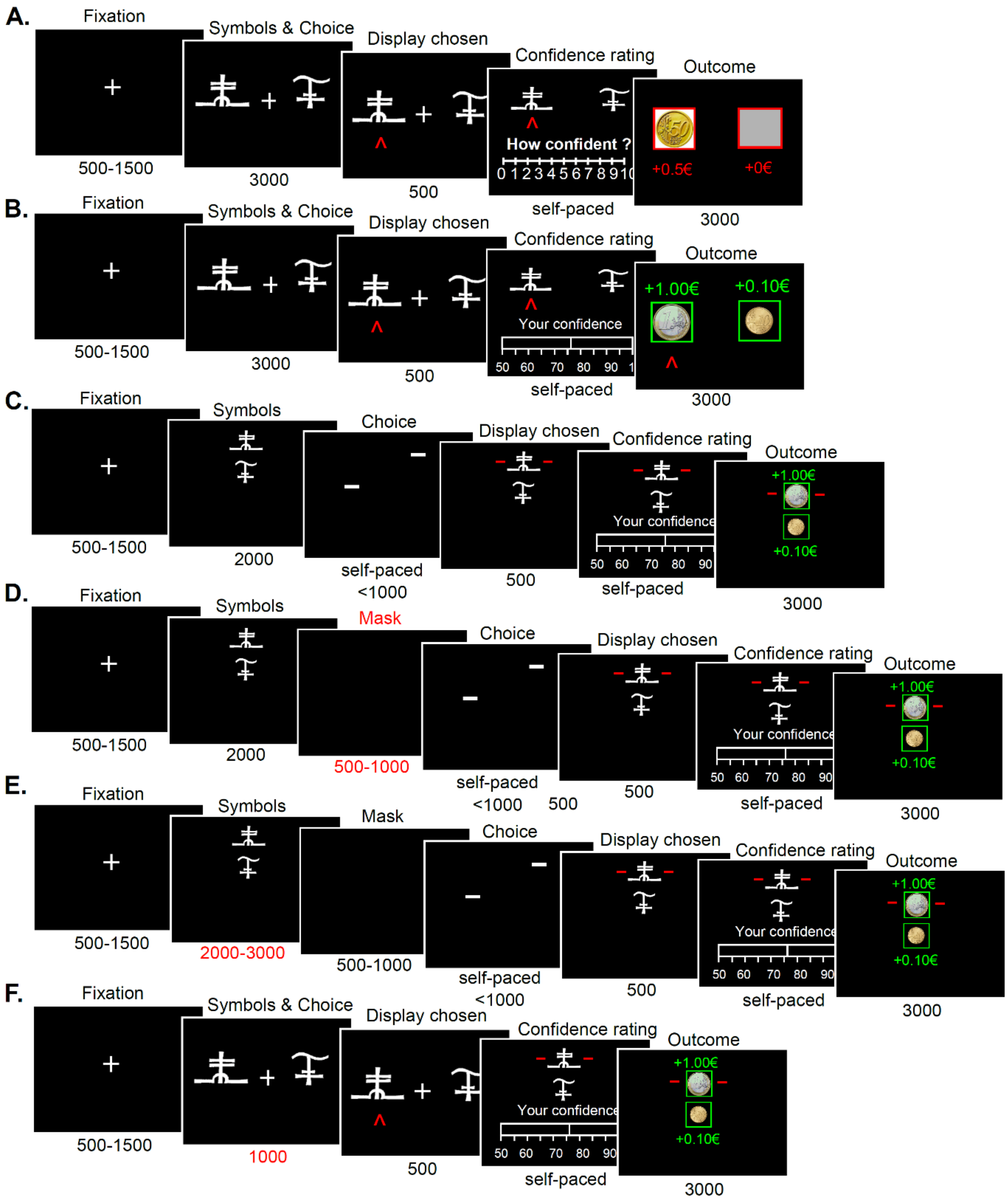


**Figure S. 1. Experimental paradigms.**

**(A-F)** Behavioral tasks for Experiments 1-6. Successive screens displayed in one trial are shown from left to right with durations in ms. All tasks are based on the same principle: after a fixation cross, participants are presented with a couple of abstract symbols displayed on a computer screen and have to choose between them. They are thereafter asked to report their confidence in their choice on a numerical scale. Outcome associated with the chosen symbol is revealed, sometimes paired with the outcome associated with the unchosen symbol -depending on the condition. Tasks specificities are as follow: **(A)** Experiment 1: symbols are displayed on the left and right sides of the screen. Confidence is reported on a 0-10 Likert scale non-incentivized. **(B)** Experiment 2: similar to experiment 1, except that confidence is reported on a 50-100% rating scale and incentivized. **(C)** Experiment 3: similar to Experiment 2, except that options are displayed on a vertical axis. Besides, the response mapping (how the left vs right arrow map to the upper vs lower symbol) is only presented after the symbol display, and the response has to be given within one second of the response mapping screen onset. **(D)** Experiment 4: similar to experiment 3, except that a short empty screen is used as a mask, between the symbol display and the response mapping. **(E)** Experiment 5: similar to experiment 4, except that a jitter is introduced in the symbol presentation. **(F)** Experiment 6: similar to experiment 2, except that a shorter duration is allowed from the symbol presentation to the choice

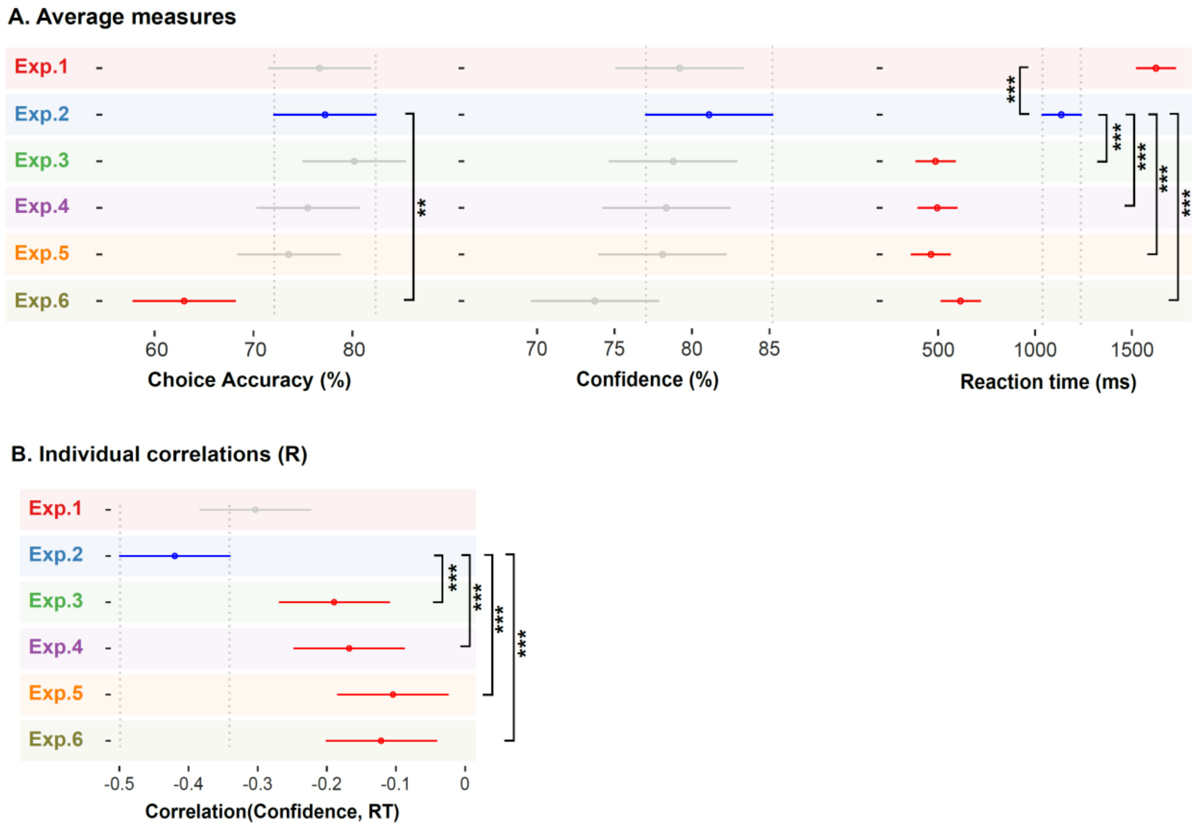


**Figure S. 2. Post-hoc analysis results.** The post-hoc tests were performed using the multcompare function in Matlab, that implements Tukey's honestly significant difference criterion to deal with multiple comparisons. Means of Choice Accuracy, Confidence and RT for each experiment are represented by a circle. The 95% confidence interval (CI) is represented by a line extending out of the circle. The post-hoc analysis compared each experiment with experiment 2 (which served as a basis to design experiments 3-6). (A) Left: **Accuracy**. Middle: **Confidence**. Right: **Response times**. (B) **Correlation between confidence and RT**. Experiment 2 is represented by a blue dot ± 95% CI error bar. Experiments represented with a red (resp. grey) dot ± 95% CI error bars were significantly (resp. not significantly) different from Experiment 2.
~ $P<0.1$; * $P<0.05$; ** $P<0.01$; *** $P<0.001$

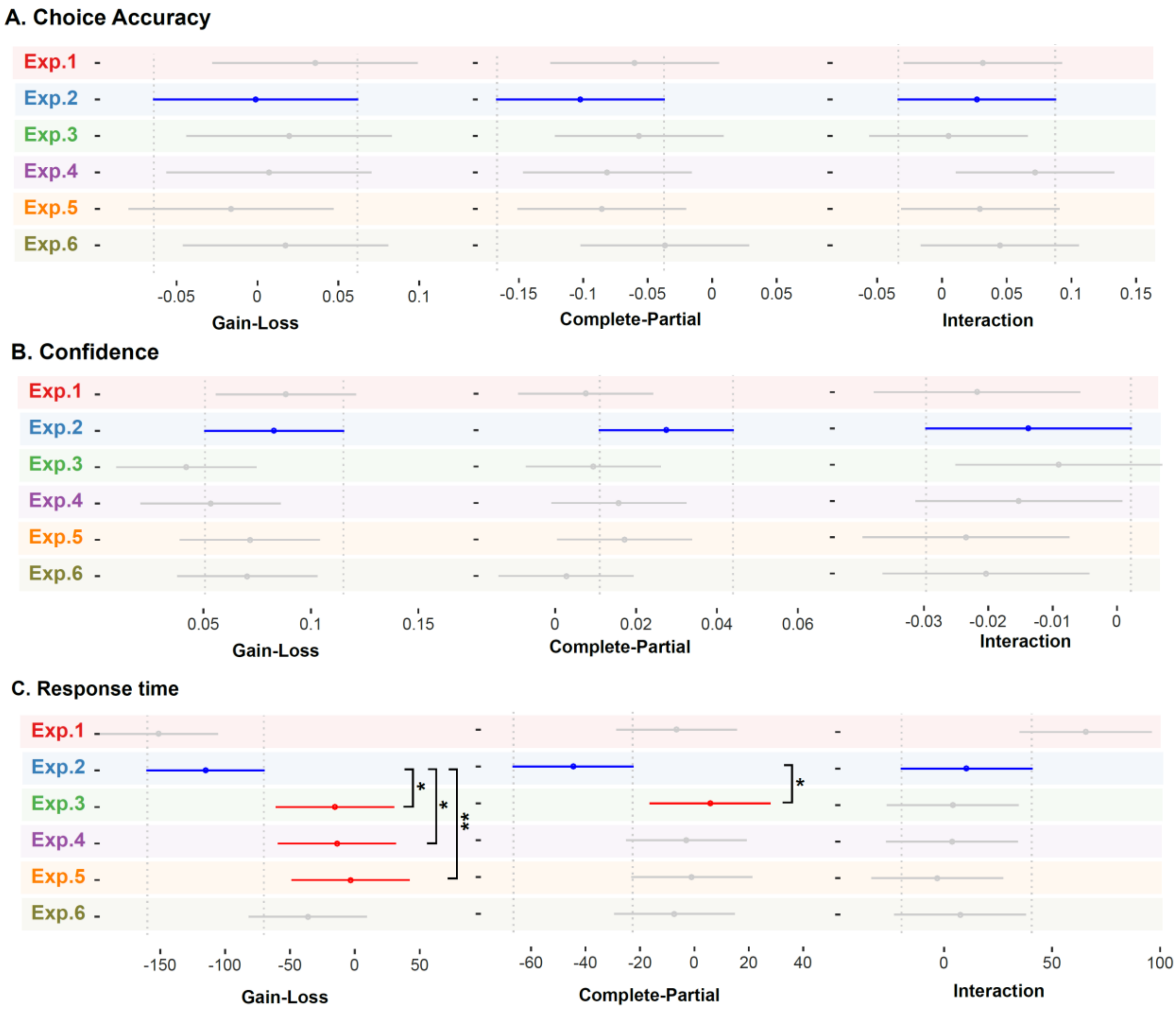


**Figure S. 3. Post-hoc analysis with multiple comparison test.** The post-hoc tests were performed with the multcompare function in Matlab, and implement Tukey's honestly significant difference criterion to deal with multiple comparisons. The mean effects of the experimental treatments (Valence, Information and their interaction) for each experiment are represented by a circle. The 95% confidence interval (CI) is represented by a line extending out of the circle. The post-hoc analysis compared each experiment with experiment 2 (whose experimental design served as a basis for experiments 3-6). (A) Effects of Valence (gain-loss), Information (complete – partial) and their Interaction on Accuracy. (B) Effects of Valence (gain-loss), Information (complete – partial) and their Interaction on Confidence. (C) Effects of Valence (gain-loss), Information (complete – partial) and their Interaction on RT.

Experiment 2 is represented with a blue dot ± 95% CI error bar. Experiments represented with a red (resp. grey) dot ± 95% CI error bars were significantly (resp. not significantly) different from Experiment 2.

~ P<0.1; * P<0.05; ** P<0.01; *** P<0.001
